# Investigating the effects of protons versus x-rays on radiation-induced lymphopenia after brain irradiation

**DOI:** 10.1101/2024.03.02.583088

**Authors:** Julie Coupey, Thao Nguyen Pham, Jérôme Toutain, Viktoriia Ivanova, Erika Hue, Charly Helaine, Ali Ismail, Romaric Saulnier, Gael Simonin, Marc Rousseau, Cyril Moignier, Juliette Thariat, Samuel Valable

**Affiliations:** Normandie Univ., UNICAEN, CNRS, ISTCT, GIP CYCERON, 14074 Caen, France; Laboratoire de Physique Corpusculaire, UMR 6534 IN2P3/ENSICAEN – Normandie Université, France; LABEO, Normandie Univ., BIOTARGEN, UR 7450, 14280 Saint-Contest, France; UAR3408/US50, UNICAEN-CNRRS-INSERM-CEA, Cyceron, GIP CYCERON, 14074 Caen, France; Université de Strasbourg, CNRS, IPHC UMR 7178, Strasbourg, F-67000, France; Medical physics department, Centre François Baclesse, Caen, France; Department of Radiation Oncology, Centre François Baclesse, Caen, France

**Keywords:** Brain radiotherapy, x-ray irradiation, proton therapy, inflammation, lymphopenia

## Abstract

**Background:** Conventional x-ray-based radiotherapy is a standard treatment for patients with brain tumors. However, is associated with systemic effects like lymphopenia that correlates with poor prognosis. Proton therapy has emerged as a new radiation strategy, given that the lower entry dose and absence of exit dose can be exploited to spare healthy brain tissues and reduce side-effects caused by systemic inflammation. We evaluated if brain irradiation with protons could spare circulating leukocytes along with other variables in rodent models.

**Methods:** Tumor-free C57BL/6 mice were irradiated with a total dose of 20Gy in 2.5Gy twice-daily sessions over four consecutive days with either x-rays or protons. Groups of mice were defined according to irradiation volume (whole-brain or hemisphere) and dose rate (1 or 2Gy/min). Blood was withdrawn at various time points and circulating lymphoid, with myeloid subpopulations analyzed using flow cytometry. Brain tissue histochemical analyses were performed late after irradiation.

**Results:** Blood sampling showed severe and acute radiation-induced lymphopenia after x-rays, with marked depletion of 50% CD4^+^ and CD8^+^, as well as B and NK cells. With protons, the decrease was 20% on average for whole-brain irradiations, suggesting a conservative effect on circulating lymphocytes. The data showed no effect in CD11b^+^ myeloid cells for both x-rays and protons. Histological analyses revealed a more intense expression level of CD68 and Iba1 immunostaining after x-ray irradiation. GFAP staining was well detected after both beams.

**Conclusion:** Proton therapy for brain tumors differs from photon therapy in terms of its effects on circulating cells and tissues.

**Key points:**

1. X-ray brain irradiation induced an acute severe lymphopenia, with a reduction of at least 50% lymphocytes. The whole-brain irradiation caused a more pronounced decrease in lymphocytes than hemisphere irradiation. Proton brain irradiation exhibited a conservative effect on circulating leukocytes.
2. X-ray irradiation-induced lymphopenia is followed by a recovery of all lymphocyte subpopulations to control levels. However, this recovery is longer for CD3^+^ lymphocytes, and B and NK cells, depending on irradiation modalities.
3. Long-term brain tissue histochemical analyses demonstrated differences between the two beams, consisting of a macrophage/microglial activation seen mostly after x-rays while an astrocyte reaction was seen after brain exposure to the two beams. These differences may explain the disparities observed in leukocytes, thereby favoring a specific biological reaction between the brain and blood.

**Importance of the Study:** Our study demonstrated that while whole-brain or hemispheric irradiation with x-rays resulted in lymphopenia, proton brain irradiation exhibited a conservative effect on circulating lymphocytes, which was paralleled by a less intense brain tissue reaction.

## INTRODUCTION

The immune system is composed of various cells commonly divided into lymphoid and myeloid lineages that ensure the immune response, along with the soluble molecules they secrete, such as cytokines, chemokines, and inflammation mediators.

Lymphocytes form a phenotypically and functionally heterogeneous immune population^1,2^. They comprise three main cell types; including B and T lymphocytes, as well as natural killer (NK) cells. NK are involved in innate immunity. These cells recognize and kill virus-infected cells, as well as some tumor cells, by triggering their apoptosis. B and T cells are responsible for the adaptive immunity. These cells differentiate from hematopoietic stem cells and mature within the primary lymphoid organs (bone marrow, thymus, and fetal liver). They are present in the bloodstream and perform their functions within the secondary lymphoid organs (spleen, lymph nodes, and mucosa-associated lymphoid tissues)^3^. T cells are divided into several lineages, including cytotoxic CD8^+^ T cells, conventional CD4^+^ helper-auxiliary T cells, and regulatory T cells (Treg). Myeloid cells contribute to the innate immune response. They mainly comprise two cell types, consisting of neutrophils designed for an efficient defense barrier against pathogens^4^, as well as monocytes, which are phagocytic cells homing into tissues to differentiate into macrophages^5,6^.

Beyond their physiological functions, these cells play an essential role regarding the tumor microenvironment (TME) and can contribute to inflammation. In recent years, inflammation has gained a great interest in the field of cancer. Indeed, inflammatory and immune cells play a central role in the control of tumor growth and in treatment responses^7,8,9^. In specific conditions, such as high-grade glioma (HGG) tumors, the number of immune cells is low^10^, contributing to poor immunotherapy responses^11,12,13,14^. In these tumors, macrophages represent the main type of inflammatory cells^10,15^, and they are associated with poor prognosis^16^. The brain TME is therefore considered as immunosuppressive. Lymphopenia (<200/µL vs. 1000/µL in treatment-naïve control patients)^17^ could be either tumor-related or result from treatment effects.

Ionizing radiations, which are applied to kill tumor cells, directly act on the tumor cells’ biological functions within the TME. Radiation-induced lymphopenia (RIL) has been reported after irradiation of several cancer forms, including central nervous system (CNS) tumors. Lymphocytes are the most radiosensitive immune cells, with a D10 (dose required for a 10% survival fraction) estimated at 3Gy^18^. In glioblastoma (GBM), over 40% of patients displayed CD4^+^ lymphocyte counts below 200/mm^3^ (500/mm^3^<baseline<1500/mm^3^) after chemo-radiotherapy^19^. Furthermore, in an *in silico* study, Yovino *et al*. 2013 reported that after 30 fractions of radiotherapy (RT), 99% of circulating cells received a lethal dose^20^. RIL can persist for several months, which is associated with poor prognosis^19,21^. RIL incidence and severity depend on dosimetric factors, treatment duration, fractionation, radiation type, and baseline lymphocyte count^22,23^. In order to circumvent x-rays’ side-effects, other radiotherapy modalities including proton therapy have emerged.

Using a four-dimensional human blood flow model, Hammi *et al*. 2020 demonstrated that, after a fraction of irradiation, the irradiated blood volume was smaller when using protons than photons^22^. Similarly, in GBM patients, proton therapy significantly reduced the ‘whole-brain mean dose,’ which was paralleled by a lower RIL incidence^23^. Taken together, these data suggest that proton therapy would likely be of interest to protect lymphocytes and preserve the anti-tumoral immune response. The radio-sensitivity of lymphocytes irradiated *in vitro* with protons may not be differ (though more pronounced) from x-ray effects^24,25^. It could be mediated by changes in cells and tissues that could, in turn, modulate the RIL extent. Additionally, little is known about the mechanisms underlying differential immune cell depletion after proton or photon irradiations. No studies are currently available detailing the effects of brain irradiation on each lymphocyte and myeloid subpopulation. Similarly, whether the effects of brain proton irradiation on circulating leukocytes are ballistic, biological, or both is still unclear.

The radiation impact on the TME, including more specifically the immune response, is of current interest, given the urgent need for targeted and immuno-therapeutic cancer treatments. Herein, we have explored the presence of systemic inflammation by measuring leukocyte subpopulations in C57BL/6 mice after brain irradiation. To avoid confounding tumor-related effects, our research was performed using non-tumor-bearing, immunocompetent mouse models.

First, changes in circulating myeloid and lymphoid counts were measured using flow cytometry at various time points following x-ray or proton brain irradiations. Radiotherapy volume (whole brain or hemisphere) and dose rate (1 or 2Gy/min) parameters were also used as variables. In parallel, the brain tissue response was evaluated by means of immunohistochemistry. In addition, lymphocytes from mice were also irradiated using either x-rays or protons in order to assess *in vitro* their direct radio-sensitivity.

## MATERIALS & METHODS

### Animal models

Seven-week-old female C57BL/6 mice (16-20g, *Janvier labs*) were employed. After irradiation, hydrating jelly (DietGel Recovery, *HOO7-72065*, SSNIFF Spezialdiäten GmbH) was added to the cage along with moistened kibble, after which the mice were returned to the animal facility and routinely maintained. Weight was monitored three times a week. Animal investigations were performed in accordance with the current European regulations, with permission of the regional committee on animal ethics CENOMEXA for the experiments carried out in Caen (#27343) and CEEA035 for those performed in Strasbourg (#27413).

### X-ray and proton irradiations

Both x-ray and proton irradiation sessions delivered a physical dose of 2.5Gy (relative biological effectiveness, RBE). Mice were irradiated under anesthesia (1L/min with 5% isoflurane for induction and 2% for maintenance in 70% NO_2_/30% O_2_), twice daily for four consecutive days, up to a total dose of 20Gy. Four experimental groups were constituted for both beams: hemispheric 2Gy/min; whole-brain 2Gy/min; hemispheric 1Gy/min; whole-brain 1Gy/min (**supplementary Figure 1**). Both x-ray and proton irradiation protocols involved 40 irradiated and 8 control anesthetized mice (CT).

X-ray irradiations were performed on a small animal irradiator (XRad-225Cx, Cyceron, Caen, France). An 8mm square collimator allowed for whole-brain or left hemisphere irradiations after identification by the inboard scanner in order to spare the olfactory bulbs and brainstem. After a “Scout” control with Al filter (80keV/0.5mA), the treatment using a Cu filter (225keV/13mA) was applied.

Proton irradiations were carried out at the PRECy platform (CYRCé cyclotron, Strasbourg, France). These irradiation sessions were performed with square or rectangular collimators enabling whole-brain or hemispheric irradiations. The energy of the proton beam was 25MeV to accurately target the brain area, and not beyond (**supplementary Figure 1**). Anesthetized mice were placed in a neck cradle. A laser pointer was employed to calibrate the irradiation brain center in a way to spare the olfactory bulbs, skull base nodes and vessels, and the brainstem, as well.

### Blood collection

For each mouse, 110µL of peripheral blood was collected submandibularly before (D_0_), during (D_3_) and after (D_7_, D_14_, D_21_, and D_28_) radiotherapy using a sterile lancet (*GR-4MM*, BiosebLab).

### Flow cytometry

#### Blood cell labeling

Blood sample (20µL) were transferred onto 96-wells V-bottom plates, treated twice with 1X red blood cell lysis buffer (*00433357*, Thermo Fisher) at room temperature (RT), then washed with 50µL of Fluorescence Activated Cell Sorting (FACS) buffer (PBS 1X/2.92% EDTA/0.2% BSA/0.2% sodium azide). Fcγ receptors were blocked using anti-CD16 and anti-CD32 antibodies (1/1000, TruStain FcX^TM^, *101320*, Biolegend) in FACS buffer for 15min at RT. Cells were then incubated for 30min in 50µL extracellular mix, containing the specific antibodies (**supplementary Table 1**); cells were subsequently incubated with an aqua vivid live/dead cell marker for 30min at RT (1/1000 in PBS, *11500597*, Thermo Fisher), fixed (Fixation/Perm Diluent, *2254166* with permeabilization concentrate, *2220751*, Thermo Fisher) and left for 30min at RT to be re-suspended in 100µL of FACS buffer. Samples were stored at 4°C and protected from light. Then, for the intracellular labeling, cells were rinsed with 50µL of 1X perm buffer (*00833356*, Thermo Fisher) and incubated for 1h at RT in the intracellular mix containing specific antibodies (**supplementary Table 1**). After a washing step with perm buffer, cells were re-suspended in 100µL of FACS buffer and stored in the dark at 4°C. Prior to the cytometer run, 10µL of counting beads (*11570066*, Thermo Fisher) were deposited in each well.

#### Raw data acquisition

Three flow cytometry panels have previously been designed to detect each myeloid and lymphocyte subtype. Cells were analyzed on a Cytoflex S^®^ (Beckman Coulter, Flow cytometry accommodation, Normandie equine Vallee platform, LABEO BIOTARGEN, Saint-Contest, France).

#### Cytometry data analysis

Cytometry data were processed using the Kaluza^®^ software (Beckman). Cells were initially selected according to their size (forward scatter, FSC) and granularity (side scatter, SSC). Doublets were excluded. For lymphocyte populations (CD45^+^CD3^+^), cytotoxic (CD8^+^), conventional (CD4^+^) and regulatory (CD4^+^FoxP3^+^) T cells, as well as B (CD45^+^CD3^-^B220^+^) and NK (CD45^+^CD3^-^NK1.1^+^) cells were evaluated. For myeloid populations (CD45^+^CD11b^+^), monocytes (SiglecF^-^Ly6C^+^) and neutrophils (SiglecF^-^Ly6G^+^) were evaluated. For each well, cell counts for each population and number of beads were recorded. The number of cells/µL was calculated and normalized by the number of counting beads recorded.

### Imaging

Animals were anesthetized, and brain and whole-body magnetic resonance imaging (MRI) were performed at 7 Tesla (Bruker, Cyceron) at D35. T2-weighted sequences were employed (RARE=8, TR/TE=5000/56msec, number of average=2, FOV=20*20mm, spatial resolution=0.078*0.078mm, 20 slices of 0.5mm thickness, and acquisition time=4min).

### Immunohistochemistry

At D_45_, mice were deeply anesthetized and transcardially perfused using a heparin/saline solution over 2 hours (h) after buprenorphine (*54000561*, Buprécare^®^) injection (0.05mg/Kg). The brain and spleen were removed and fixed in isopentane. Before freezing, the spleen was placed in RPMI medium (*RO883*, Sigma-Aldrich) and weighted. The 20µm thick cryostat brain sections were performed and stored at -80°C. Slides were rehydrated and blocked with PBS/0.1% Tween/0.5% Triton/3% BSA for 2 hours and incubated overnight with primary antibodies in PBS/0.1% Tween/0.5% Triton/1% BSA at 4°C. Sections were then incubated with fluorochrome-conjugated secondary antibodies and Hoechst 33342 (*14533*, Sigma-Aldrich) in PBS/0.1% Tween/0.5% Triton/1% BSA.

### T cell isolation from the spleen

#### Sample preparation

The EasySep^TM^ Mouse T Cell Isolation Kit (*19851*, StemCell) was used to isolate lymphocytes from single-cell splenocyte suspensions by negative selection. Spleens were withdrawn on 7-week-old female Swiss mice (22-25g, CURB@CYCERON) and manually dissociated with 10mL of PBS/2% Fetal Calf Serum (FCS). Aggregates and debris were removed by passing the suspension through a 70µm mesh strainer (*27260*, StemCell). The obtained solution was centrifuged at 300g for 5 minutes (min) and re-suspended after counting at 1.10^8^ cells/mL in PBS/2% serum.

#### T cell isolation

Overall, 2mL of the prepared sample were transferred into a 5mL polystyrene round-bottom tube. Cells were stained with 50µL/mL of biotinylated antibodies and 75µL/mL of streptavidin-coated magnetic particles. Undesired cells were separated using an EasySep^TM^ magnet (*18000*, StemCell) by placing the tube into the magnet at RT for 2.5 min. Lymphocytes were poured off into a new tube, counted on a Malassez counting chamber; RPMI medium was then added to obtain target cell seeding concentration. Overall, 20U/mL of IL-2 (*21212*, Peprotech) and 1µg/mL of anti-CD28 (*102116*, Biolegend) were added to the isolated lymphocyte suspension before seeding at 1mL/well in a 24-well plates coated with 1µg/mL of anti-CD3ε (*100340*, Biolegend) 24 h before the procedure.

### Cell irradiation

Overall, 48 h after seeding, the lymphocytes were irradiated using either x-rays (FAXITRON, Cyceron, Caen, France) or protons from an IBA ProteusOne at the Normandy proton therapy center (CYCLHAD, Caen, France) in cell culture plates 48 h after cell seeding. Six wells were seeded (1.10^6^ cells/1.5mL) per dose condition to obtain duplicates for each counting time point (24 h, 48 h, and 72 h post-irradiation). X-ray exposure was performed at 2Gy/min (130keV/5mA/Cu filter). For protons, the LET was set at 4.6keV/µm. For both beams, cells were exposed at RT at 0.5, 1, 2, or 4Gy, after which they were maintained in culture. At each time point, the supernatant and proteins were collected for further analysis. The experiments were run three times, and the survival fraction at 2Gy was extracted using the *CS-Cal* software for clonogenic survival calculation.

### Statistical analyses

*In vivo* data were presented as mean ± standard deviation (SD). Statistical analyses were performed using GraphPad Prism 9.0 (San Diego, USA). Significances were calculated by means of non-parametric Kruskal-Wallis tests for multiple comparisons. Data were presented according to changes versus control conditions (CT) in percentages. Statistical significance was achieved when *p* <0.05 (*); *p* <0.005 (**); *p* <0.001 (***), otherwise it was not significant.

## RESULTS

### Brain irradiation using x-rays induced an acute decrease in CD45^+^ leukocytes unlike with protons

We first assessed the effects of x-rays and protons on total CD45^+^ leukocytes at D_2_ (*i.e*., upon irradiation). Plots in **Fig. 1A** clearly showed a decrease in the CD45^+^ cell amounts in comparison with controls (CT) in response to x-rays, whereas no visible effect was noted with protons. The quantification confirmed this acute effect after x-rays at D_2_, regardless of dose rate or irradiation volume. The CD45^+^ cells dropped by approximately 50% after irradiation at 1Gy/min (*p*=0.0180) for the H-irradiated group and at 2Gy/min (*p*=0.0427) for WB-irradiated group, in comparison with CT (**Fig. 1B**, *left*). No change occurred within the irradiated and CT groups after proton treatment at D_2_, irrespective of dose rate or irradiation volume (**Fig.1B**, *right*). A slight but not significant decrease in CD45^+^ leukocytes was noticed for WB-irradiated groups compared with both H-irradiated and CT groups, after proton irradiation. At D_28_, no residual difference was observed between the irradiated and CT groups after x-ray therapy (**Fig.1C**, *left*). Similarly, after proton therapy, all irradiated groups remained at CT levels (**Fig.1C**, *right*).

**Figure 1.**
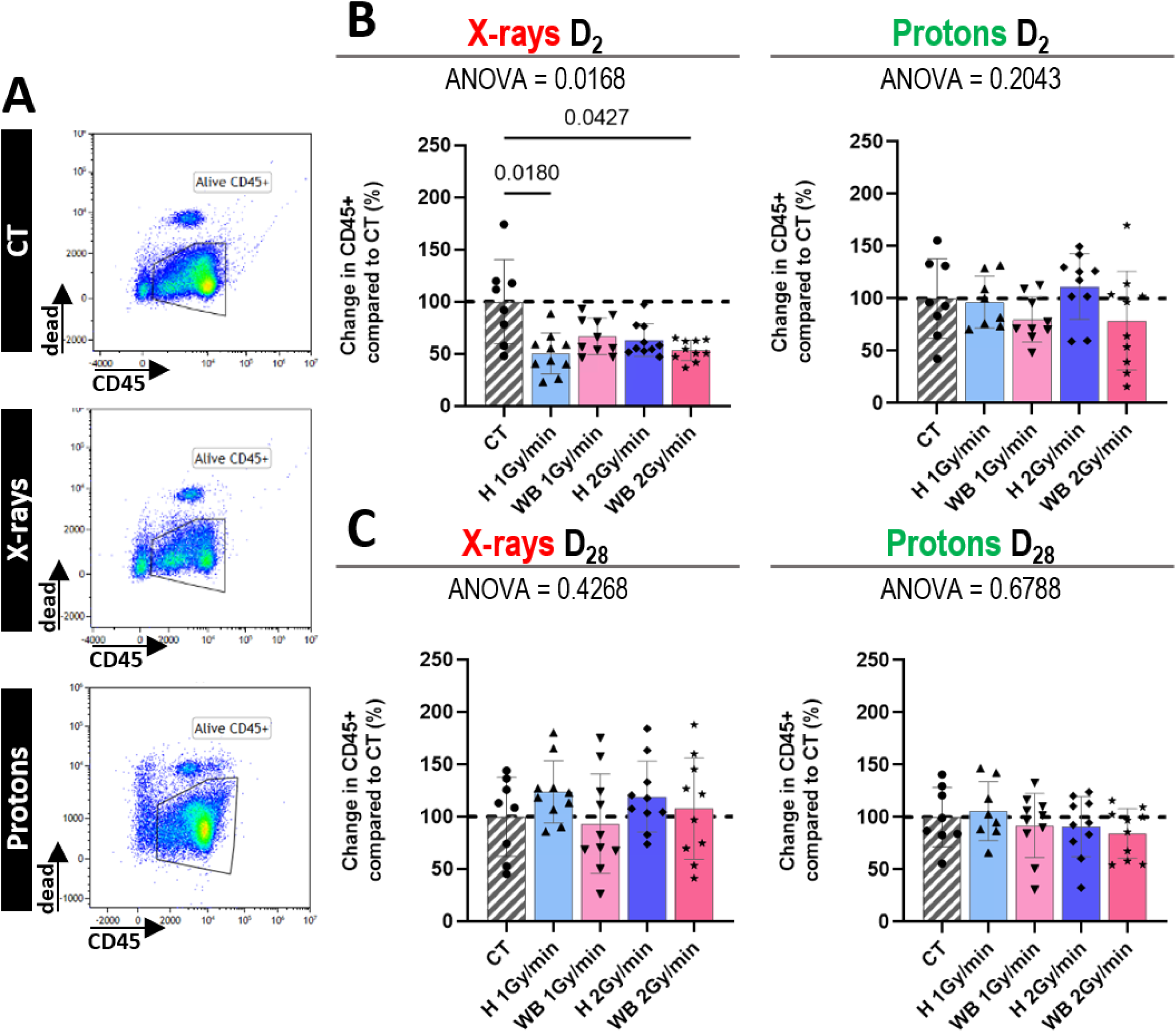
Change in CD45^+^ leukocytes after conventional x-ray or proton irradiation. The histograms represent changes in the total number of CD45^+^ leukocytes at Day 2 (D_2_) **[B]** or Day 28 (D_28_) **[C]** after x-ray or proton exposure. All data were represented as the percentage change in irradiated groups (n=10 per groups) compared with the control group (CT) (n=8). Data were expressed as mean ± standard deviation. *P*-values were determined using a one-way ANOVA test, along with a multiple comparison Kruskal-Wallis test. Dot plots on the left illustrate the variation of cells at D_2_ **[A]** compared with CT for WB groups following x-ray or proton exposure at 2Gy/min.

### Brain irradiation with x-rays induced an acute decrease in CD3^+^, CD8^+^, B, and NK cells unlike with protons

At D_2_, results showed a significant decrease in CD3^+^, CD8^+^, and B and NK cells. (**Fig.2A**). This effect was particularly pronounced in WB 2Gy/min-irradiated groups (p=0.0011 for CD3^+^, p=0.0001 for CD8^+^) compared with CT. There was a significant difference in CD3^+^ (p=0.0052 and p=0.0575) counts between CT and irradiated groups at 1Gy/min regardless of the irradiation volume. The effect was more pronounced for CD8^+^ cells in all irradiated groups (p=0.0018 for H at 1Gy/min; p=0.0160 for WB at 1Gy/min; p=0.0474 for H 2Gy/min; p=0.0001 for WB 2Gy/min). Of note, a non-significant decrease in CD4^+^cells was observed.

**Figure 2.**
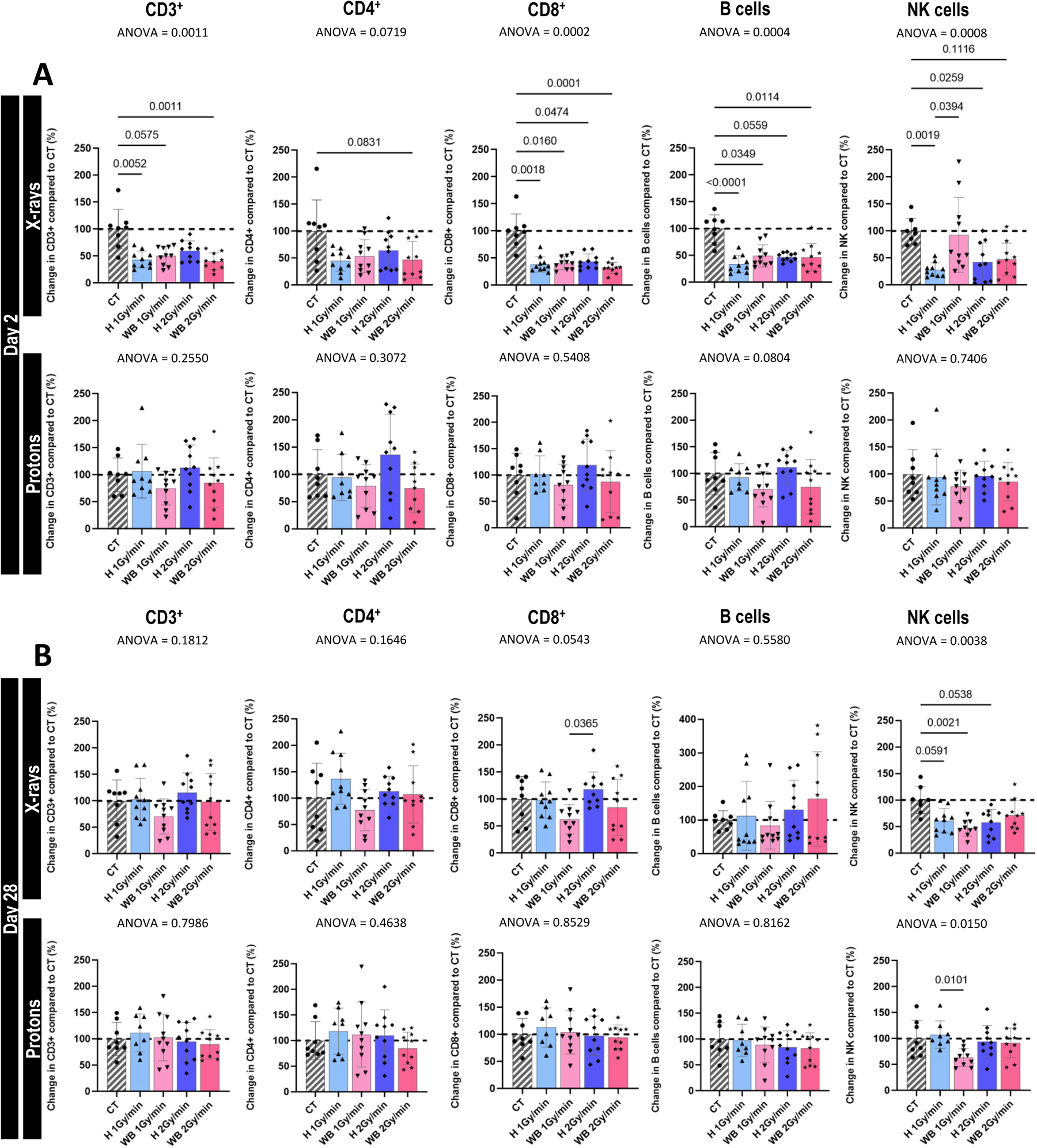
Changes in lymphoid populations early [A] or late [B] after brain x-ray or proton irradiation. The histograms represent changes in the total number of CD3^+^, CD4^+^, and CD8^+^ T subpopulations, B and natural killer (NK) cells at Day 2 (D2) **[A]** or Day 28 (D28) **[B]** after x-ray or proton exposure. All data were represented as the percentage change in irradiated groups (n=10 per groups) compared with the control group (CT) (n=8). Data were expressed as mean ± standard deviation. *P*-values were determined using a one-way ANOVA test, along with a multiple comparison Kruskal-Wallis test.

The B cell population dropped by more than two-thirds compared with CT at D_2_ after x-ray treatment in WB-irradiated groups regardless of the dose rate (p=0.0349 at 1Gy/min; p=0.0114 at 2Gy/min). Similar observations were made between CT and H-irradiated group at 1Gy/min (p <0.0001) (**Fig.2A**). Half of the NK cell population significantly decreased after x-ray therapy at D_2_ for the H-irradiated groups at 1Gy/min (p=0.0019) and 2Gy/min (p=0.0259). No significant differences were observed in heterogeneous WB-irradiated groups. There was no significant disparity between CT and proton-irradiated groups at D_2_, irrespective of the exposure variables (volume or dose rate). Nevertheless, there was a likely tendency for depletion in CD3^+^, CD4^+^, CD8^+^, as well as B and NK cells in WB-irradiated groups regardless of the dose rate (**Fig.2A**), with no effect observed for the other irradiated groups.

At D_28_, a recovery was almost reached for CD3^+^ and CD8^+^ in each irradiated group (**Fig.2B**). For H-irradiated groups, this recovery even went beyond the CT baseline level compared with the WB-irradiated groups at 1Gy/min where no recovery was yet reached. There was also a recovery for B cells beyond baseline levels, yet with high heterogeneity. However, the significant NK cell loss persisted compared with CT regardless of irradiation volume or dose rate conditions. At D_28_ after proton irradiation, no significant dissimilarity among the groups regarding CD3^+^, CD4^+^, CD8^+^ and B cells basal level were recorded, regardless of irradiation conditions. For NK cells, the WB-irradiated group at 1Gy/min still exhibited a cell loss of about one-third compared with CT (p=0.0101) (**Fig.2B**).

### X-ray and proton brain irradiation did not modify CD11b^+^ myeloid cell counts

Our next step consisted in evaluating the effect of x-rays or protons on CD45^+^CD11b^+^ myeloid cells, CD45^+^CD11b^+^Ly6G^+^ neutrophils and CD45^+^CD11b^+^Ly6C^+^ monocytes.

At D_2_ after x-ray exposure, there was no disparity among irradiated groups and CT (**Fig.3A**). However, there was an increase in CD11b^+^ cells (p=0.0289), and in neutrophils (p=0.0268) between WB- and H-irradiated groups at 1Gy/min but not in monocytes. After proton therapy, there was no significant difference among the groups and CT, irrespective of the irradiation conditions.

**Figure 3.**
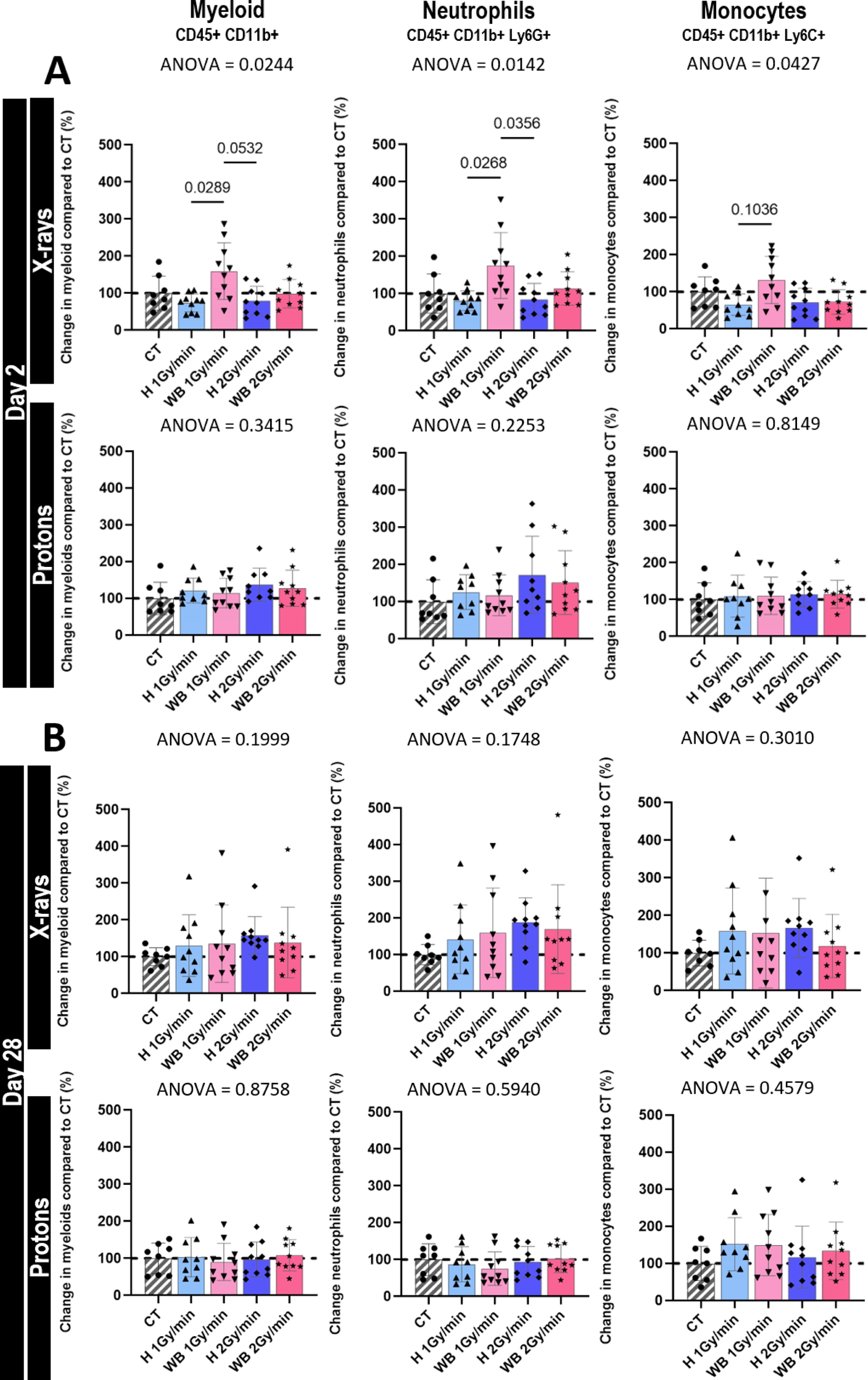
Changes in myeloid populations early [A] or late [B] after brain x-ray or proton irradiation. The histograms represent changes in the total number of CD11b^+^ myeloid, neutrophil, eosinophil and monocyte subpopulations at Day 2 (D_2_) **[A]** or Day 28 (D_28_) **[B]** after x-ray or proton exposure. All data were represented as the percentage change in irradiated groups (n=10 per groups) compared with the control group (CT) (n=8). Data were expressed as mean ± standard deviation. *P*-values were determined using a one-way ANOVA test, along with a multiple comparison Kruskal-Wallis test.

At D_28_ after x-ray treatment, there was recovery of myeloid cells, neutrophils and monocytes beyond the CT level for all irradiated groups irrespective of the irradiation conditions (**Fig.3B**). Concerning proton exposure at D_28_, no dissimilarity was observed among the irradiated groups and CT (**Fig.3B**).

### After x-ray irradiation, recovery to baseline values is cell type dependent

We then assessed the recovery kinetics between D_2_ and D_28_ for each cell subpopulation (**Fig.4A**). For CD45^+^, the decrease at D_2_ was followed by a recovery at D_14_ (**Fig.4A**, *arrow*) that went beyond the CT level for hemispheric irradiation, and stabilized until D_28_. For WB irradiation with x-rays, it reached the CT level at D_28_ whereas cell counts were stable without depletion or rebound, after proton exposure. For CD3^+^, cell counts reached CT baseline at D_21_ while for CD4^+^ it occurred at D_14_ (for hemisphere irradiations) or D_21_ (for WB irradiations). For CD8^+^, the time required to reach CT level was D_28_ for the hemispheric irradiation but remained below for the WB irradiation.

**Figure 4.**
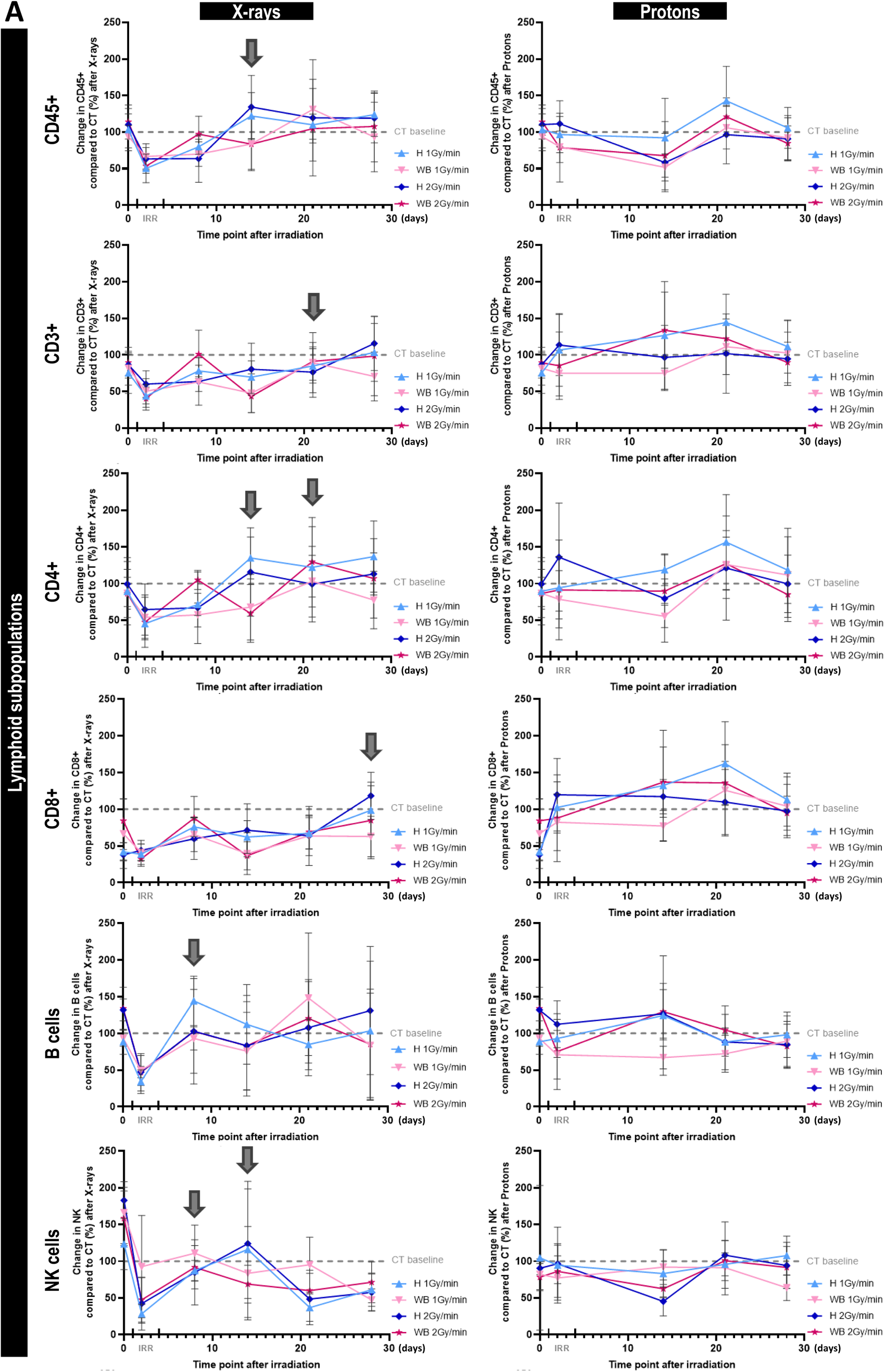
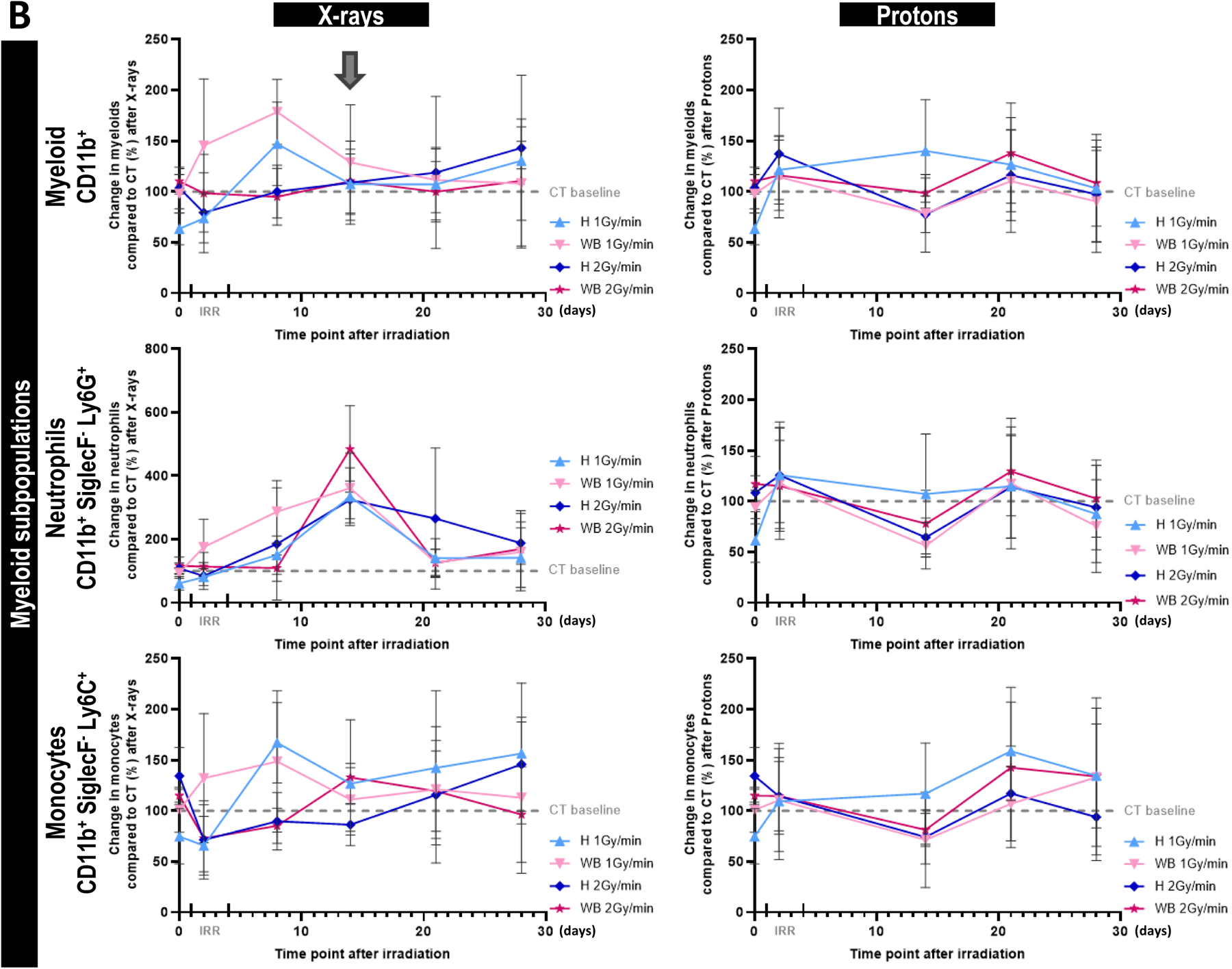
Kinetics of lymphoid and myeloid subpopulations over time after x-ray or proton brain irradiation. The curves show the evolution of lymphoid **[A]** and myeloid **[B]** subpopulation counts compared with controls (CT) over time (D_0_, D_2_, D_7_, D_14_, D_21_, and D_28_ for x-rays and D_0_, D_2_, D_14_, D_21_ and D_28_ for protons). Gray arrows correspond to the time point of recovery to baseline. The percentage evolution in cell counts from irradiated groups (n=10 per groups) compared with CT (n=8) is shown. All data were expressed as mean ± standard deviation.

Seven days were needed for B cells, while for NK cells, it was D_7_ and D_14_ respectively for H and WB irradiation. However, for x-ray treated NK cells, the recovery at D_14_ was followed by a drop at D_21_ for all groups without returning to the baseline at D_28_.

After x-ray and proton exposure, the data showed that CD11b^+^ count varied a little among groups in comparison with CT but with heterogeneity. No recovery was observed since no acute lymphopenia was detected (**Fig.4B**).

### Crosstalk between brain irradiation, blood compartments, and peripheral organs

In parallel to the follow-up on leukocyte changes, animal weight was monitored as readout of their general condition. As expected, regardless of the cohort, control mice gained weight with time (**supplementary Fig.3A**). After x-ray treatment, a significant weight loss was observed among CT and WB-irradiated groups and, H-irradiated group at 1Gy/min (**supplementary Fig.2** and **supplementary Fig.3**). After x-rays exposure, the maximum weight loss was reached at D_10_ after irradiation (on average -2.721g, **supplementary Fig.2**). After 14 days, mice regained their initial weight but remained below CT over time. After proton exposure, the weight variation was not different from CT, regardless of irradiation volume and dose rate (**supplementary Fig.3B**).

We then performed brain MRI to assess tissue changes after brain irradiation. Images showed no evidence of radio-necrosis or macro-anatomical changes after 35 days following x-ray or proton therapy (**supplementary Fig.3C**).

After euthanasia, the spleens were weighted (**supplementary Fig.3D**). Post-mortem analyses revealed a decrease in spleen weight for WB-irradiated mice compared with H-irradiated and CT. There was a significant difference (*p*=0.0260) in weight between WB-irradiated group at 2Gy/min and H-irradiated group at 1Gy/min. There was no disparity in the groups after proton-exposure.

In parallel, a whole-body MRI was performed after whole-brain or hemispherical x-ray irradiation at 2Gy/min or 1Gy/min in C57BL/6 tumor-free mice (**supplementary Fig.3E**). At D_35_, whole-body images showed a reduction in spleen size among the groups and particularly after WB irradiation (*white arrows*). At 2Gy/min the spleen volume seemed to decrease compared to 1Gy/min and CT.

### X-ray irradiation induced a more pronounced brain tissue reaction than proton irradiation

To assess the brain reaction after irradiation, we performed immunohistological analyses at D_45_. A sustained CD68^+^/Iba1^+^ labeling was observed in the left striatum after x-ray treatment on H- and WB-irradiated slices (*white arrows*) (**Fig.5A**). There was likely no disparity between dose rates after x-ray exposure, except for a greater CD68^+^ labeling on WB-irradiated sections. Almost no CD68^+^ labeling was observed following proton irradiation. There was a slight Iba1^+^ labeling, irrespective of the dose rate, after proton exposure. It is also interesting to note that on WB and H sections irradiated with x-rays at 2Gy/min, there was a CD68/Iba1 co-labeling (*white dotted circles*), especially at 1Gy/min WB-sections and at 2Gy/min slices (**Fig.5A**).

**Figure 5.**
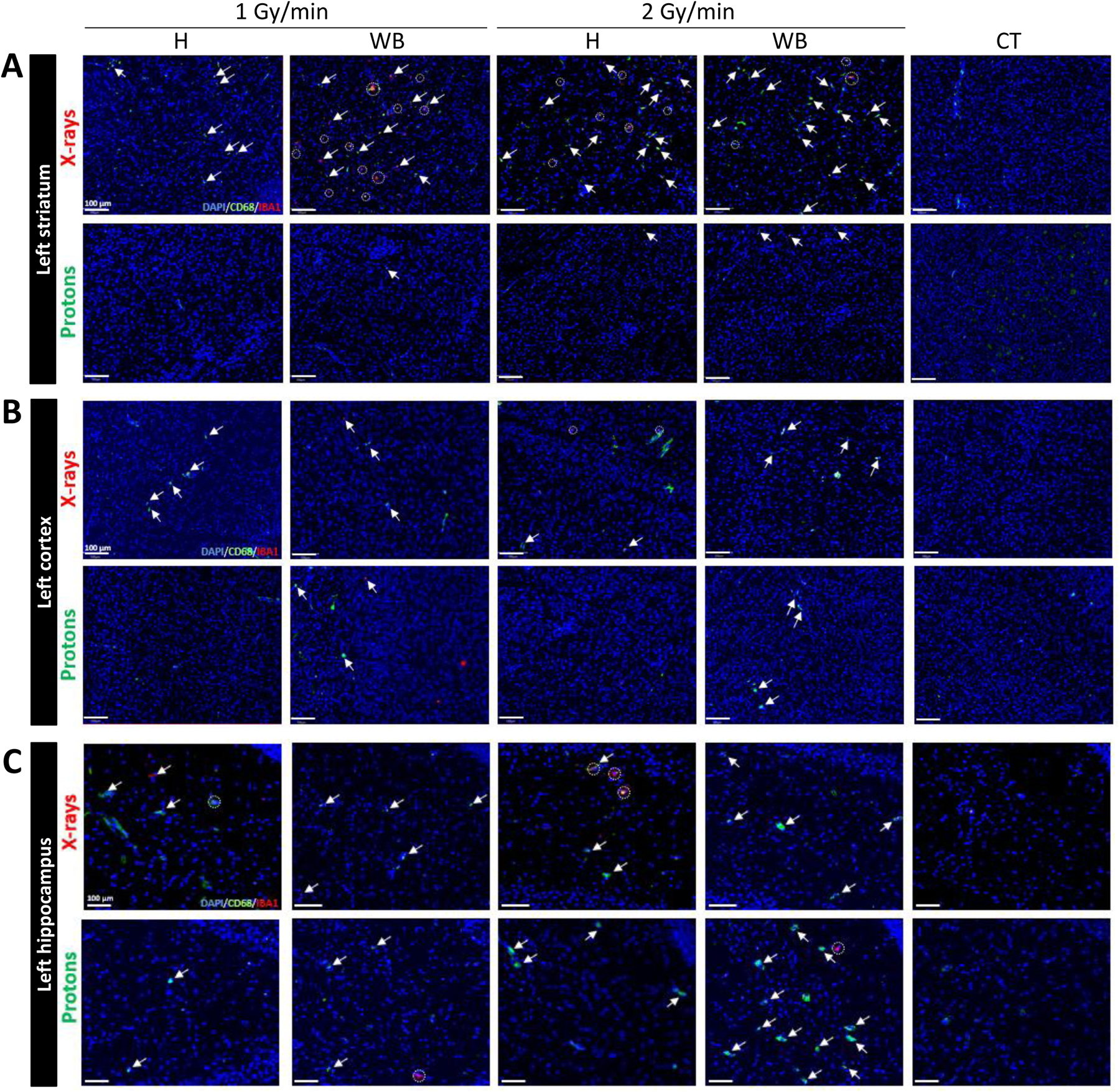
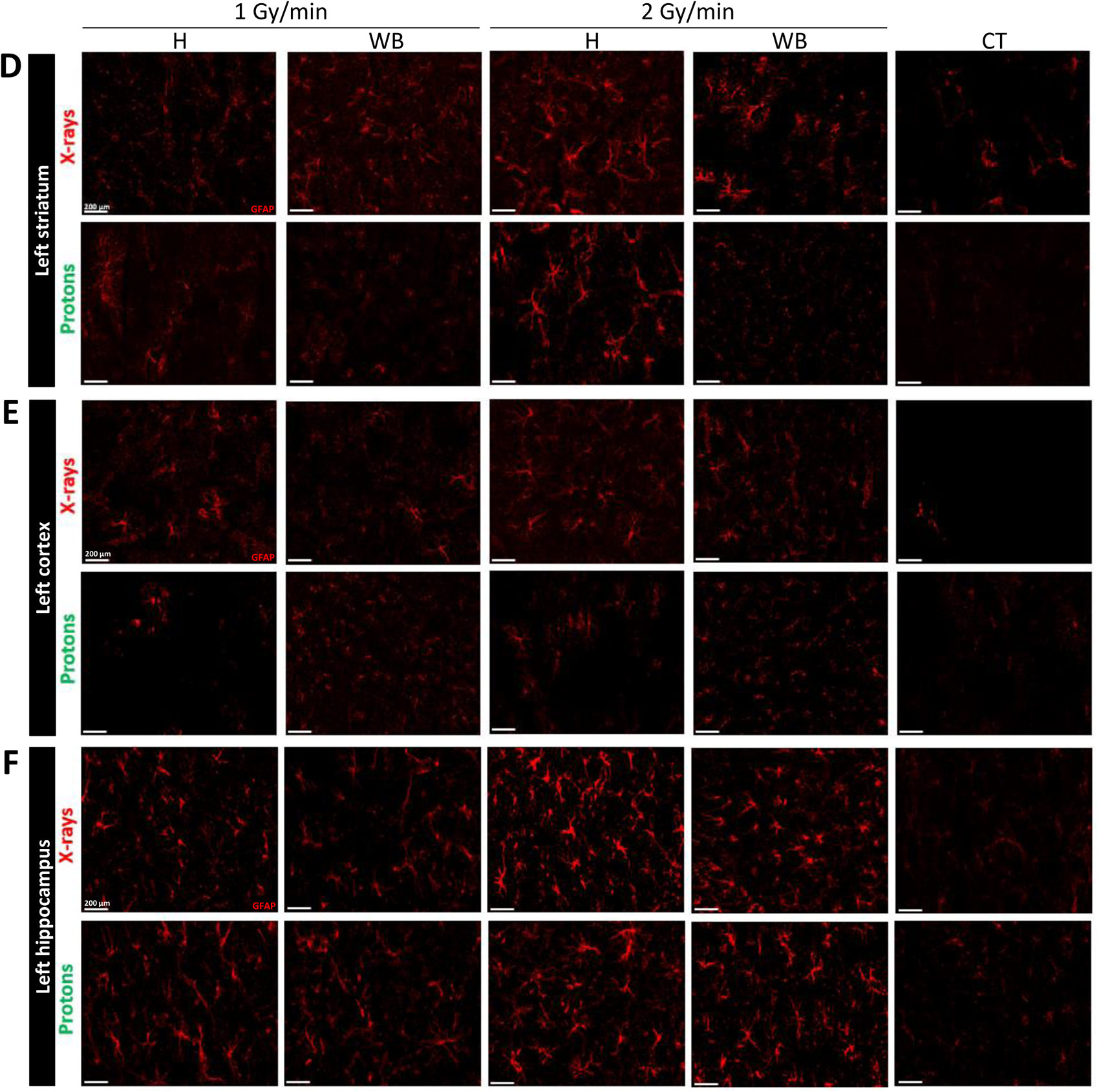
Microglia/macrophage [A, B, C] and astrocyte [D, E, F] immunostainings on brain tissues after x-ray or proton brain irradiation. [A-C] *Representative CD68 (green), Iba1 (red) and Hoechst 33342 (blue) staining of hemisphere (H) or whole-brain (WB) irradiated C57BL/6 tumor-free mice compared with controls (CT) after x-ray or proton exposure*. Scale bars = 100 µm. White arrows correspond to specific CD68^+^ or Iba1^+^ signals. Dotted white circles delimit CD68/Iba1 co-labeling. Three regions of interest (ROI) were identified on brain sections including striatum **[A]**, cortex **[B]** and hippocampus **[C]**. [D-F] *Representative GFAP (red) staining of the hemisphere after x-ray or proton irradiation of C57BL/6 tumor-free mice brains compared with controls (CT)*. Scale bars = 200 µm. Three regions of interest (ROI) were identified on sections including striatum **[D]**, cortex **[E]** and hippocampus **[F]**.

In the cortex (**Fig.5B**), only a few dots of CD68^+^ or Iba1^+^ labeling were observed. WB-irradiated sections showed positive labeling for CD68 and Iba1 after both x-ray and proton exposures. No co-labeling was observed, and it was thus difficult to distinguish specific labeling from artifacts. There was almost no labeling, except for a slight CD68^+^ staining on the WB proton-irradiated sections. The hippocampus labeling (**Fig.5C**) revealed CD68^+^ and Iba1^+^ dots, as well as co-labeling after x-ray exposure. Intense and diffuse CD68^+^ dots were also visible after WB proton irradiations at 2Gy/min.

GFAP staining was used to assess brain tissue reaction. In the left x-ray-irradiated striatum (**Fig.5D**) and cortex (**Fig.5E**), the labeling was more intense and diffuse compared with CT sections: Star-shaped astrocytes were clearly seen in WB groups regardless of dose rate and on H-irradiated sections at 2Gy/min. No disparity in GFAP labeling was observed following proton or x-ray irradiations. The labeling appeared more intense on WB section after x-ray exposure, although its intensity was also quite high on H sections irradiated by protons at 2Gy/min. The activated forms of astrocytes were clearly seen with their extensive and arborescent branches. In line with the blood samples, WB-x-ray irradiation at 2Gy/min likely exerted a greater impact, with more intense labeling, thus generating a long-term astrocytic response at the brain level after irradiation.

In the cortex, astrocytes displayed extended and arborescent extensions for all conditions following x-rays. After proton treatment, in contrast to the striatum, there was little GFAP^+^ labeling in H-irradiated sections, whereas in WB-irradiated sections, the GFAP^+^ labeling was intense. However, there were no extensions or arborescence compared with x-ray observations.

In the hippocampus (**Fig.5F**), GFAP labeling was almost similar following either x-ray or proton exposure. GFAP labeling was more diffuse and intense at 2Gy/min irradiation than at 1Gy/min due to the presence of extensions and arborescence. We also noted that the activated forms of astrocytes were particularly marked in the hippocampus following both x-ray and proton treatments.

### Isolated lymphocytes are more sensitive to protons than x-rays

We then aimed to evaluate *in vitro* the radio-sensitivity of lymphocytes to either x-rays (**Fig.6A**) and protons (**Fig.6B**) by quantifying the cell numbers at different post-irradiation times (24 h, 48 h, and 72 h) at four different doses (0, 0.5, 1, 2, and 4Gy).

**Figure 6.**
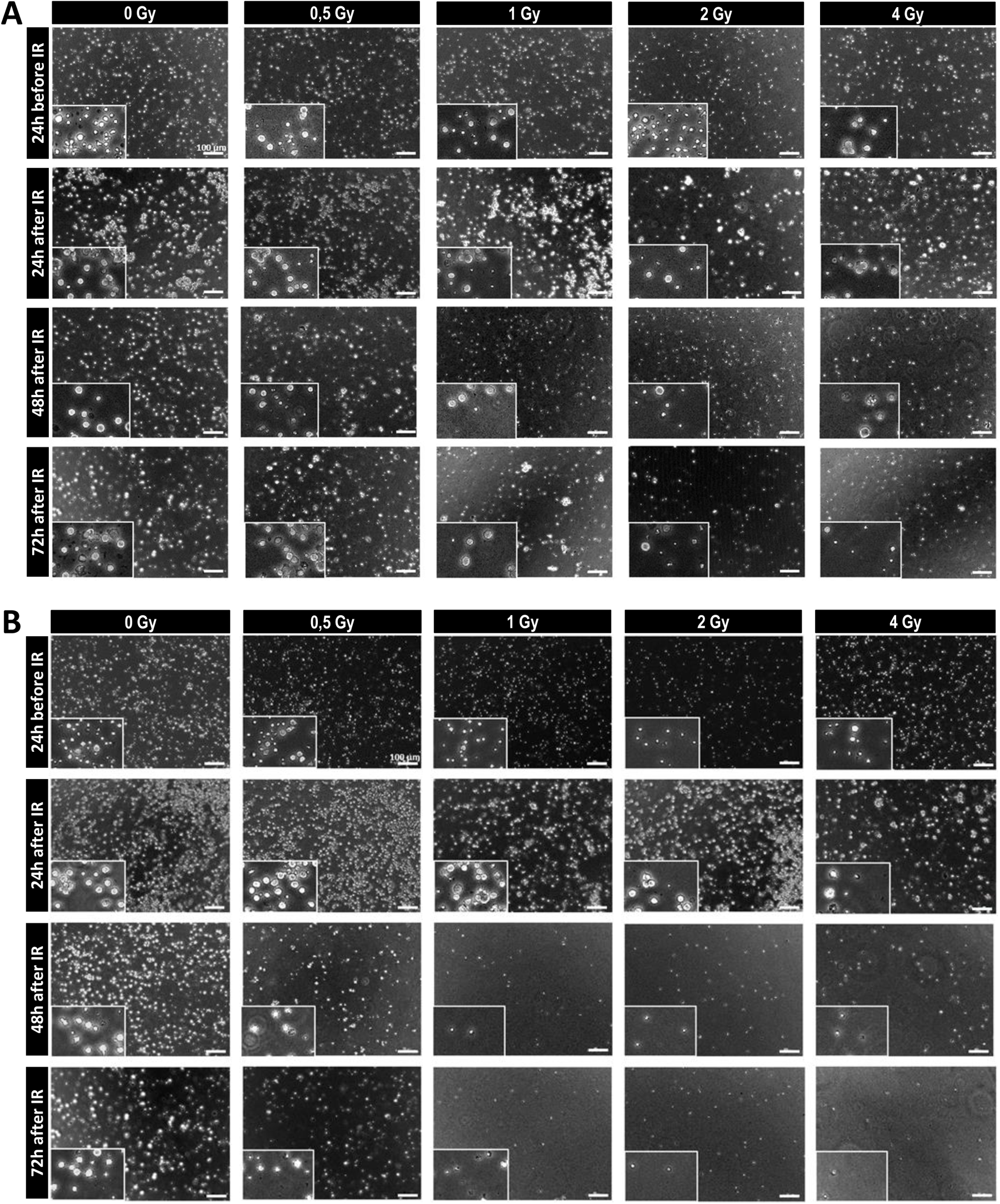
X-ray brain irradiation induced a higher reduction in shape, size, and number of T cells than proton irradiation. [A] *Representative microscopy images of isolated splenic T cells 24 h before and 24 h, 48 h and 72 h after 0, 0.5, 1, 2, and 4Gy x-ray brain irradiation (IR).* Scale bars = 100 µm. For each condition, two images were shown, namely one taken at ×10 and the other one in the bottom left corner at ×20. [B] *Representative microscopy images of isolated splenic T cells 24 h before and 24 h, 48 h and 72 h after 0, 0.5, 1, 2, and 4Gy proton brain irradiation (IR).* Scale bars = 100 µm. For each condition, two images were shown: namely one taken at ×10 and the other one in the bottom left corner at ×20.

Overall, 24 hours before x-ray irradiation, under all conditions, lymphocytes appeared round, stringent, and of a size consistent with “healthy” CD3^+^ cells according to the literature^26^. There was no dissimilarity among the doses or with CT at 24 h following x-ray irradiation, as cells remained round and stringent. Data showed a difference from 48h post-irradiation onwards. At 1, 2, and 4Gy the number of lymphocytes decreased sharply, and the live cells were smaller compared with CT. AT 72 hours post-irradiation, the number of cells was further decreased. At 2 and 4Gy, there were only scarce residuals. Therefore, number of CD3^+^ cells seemed to decrease from 48 h post-irradiation onwards for the 2 and 4Gy doses. Quantitatively, the survival fractions at 2Gy were 0.779, 0.717, and 0.580 at 24 h, 48 h, and 72 h, respectively after x-ray exposure.

The difference after 48 h was more pronounced following proton irradiation than x-rays. For the 1, 2, and 4Gy doses, the number of cells was significantly lower than in CT, as early as at 48 h. The surviving fractions at 2Gy were 0.610, 0.472, and 0.430 at 24 h, 48 h, and 72 h, respectively, after exposure.

## DISCUSSION

The present study sought to evaluate x-ray and proton effects on each lymphocyte and myeloid subpopulation after brain irradiation, as well as to decipher whether these effects were ballistic, biological, or both of them.

Our results confirmed an acute drop in circulating lymphocytes as early as D_2_ following brain x-ray irradiations corresponding to a total deposition of 10Gy. These results clearly agree with observations reported in the literature^19,20,22,27^.

More interestingly, we demonstrated a further decrease in CD3^+^, CD4^+^, CD8^+^, as well as B and NK cells. The lymphocytes radio-sensitivity is known to be high (D10=3Gy^18^). In our experimental procedure, this dose was reached, and even exceeded, on the first day of the irradiation protocol (*i.e.* 5Gy) in the brain tissues. Therefore, circulating leukocytes within the brain and head are likely to have received a significant dose. Consequently, RIL may be a direct effect of irradiation on circulating cells passing through the beam.

Proton exposure did not cause any significant effect on lymphoid or myeloid subpopulations at D_2_ compared with x-rays. However, WB proton irradiation induced a possible slight decrease in T and B lymphocytes, as well as NK cells compared with CT. Moreover, the extent of lymphopenia was independent on the irradiation dose or volume, with recovery delayed for WB relative to the hemisphere. The observed conservative effect of protons is in line with data reported in clinical studies^23^, which could be related to a smaller irradiation volume. Indeed, in our experimental set-up, protons likely spare the lower part of the brain where large arteries are located.

We recently reviewed parameters for further understanding the RIL mechanisms, based on biological theories in combination with mathematical modeling^28^. This review provided quantitative estimates of lymphocyte migration and residence times in major organs consisting of 2 h in the spleen, 10 h in lymph nodes, and less than 1 min in the lungs and liver. Only 5% of lymphocytes are present in the circulating blood compartment at all times^28^. The lymphocyte kinetics simulation in blood revealed that they recovered more than 80% of their initial level in less than 200 min after a sudden drop. These data are inconsistent with the hypothesis suggesting that leukocytes are directly irradiated through the beam. It can be assumed that there is an interaction between irradiated brain tissue, circulating blood compartment, and peripheral hematopoietic organs regulating circulating leukocyte levels.

In addition to blood parameter changes, the weight of x-ray irradiated mice significantly decreased compared with the proton-irradiated ones or CT. This could be attributed to dose deposition in salivary glands and face muscles (masticators). It is known that during radiotherapy for head and neck cancers, salivary glands are unavoidably co-irradiated causing hyposalivation and xerostomia with hard consequences for patient quality of life^29^. On the contrary, in our protocol, protons spared all sensitive organs, explaining the absence of weight loss. Interestingly, weight loss was not observed for hemispheric x-ray irradiations, whereas lymphopenia was still observed, indicating that these two parameters are not correlated.

Several days post-irradiation, brain imaging showed no macro-anatomical change in irradiated mice brain and no appearance of radionecrosis compared with CT. This is in line with literature previous results showing that 20Gy brain irradiation did not cause radionecrosis in the rodents^30^.

However, immunohistochemistry revealed a brain tissue reaction to x-rays via CD68, Iba1, and GFAP staining reflecting macrophagic/microglial and astrocyte activation. Suckert *et al*. 2021 showed that gliosis, proliferation, and astrocyte reaction occurred in the proton irradiation field, but with higher doses than ours (>40Gy)^31^. In contrast, in our case, proton exposure promoted astrocyte activation, whereas no staining was observed for CD68 and Iba1. The differences between the two beams may account for the disparities observed in leukocytes, favoring one biological reaction between the brain and blood, as discussed in the 2020 review by Cesaire *et al*^32^. We are currently further investigating this link via histological analyses and quantification of cytokines and chemokines from irradiated mice plasma samples.

In addition to immune cells in the blood, the impact on peripheral hematopoietic organs was examined. Mice’s whole-body magnetic resonance imaging MRI images showed a decrease in spleen size between H and WB following x-ray irradiation, especially at 2Gy/min compared with 1Gy/min and CT. Post-perfusion weighing revealed a decrease in spleen weight of x-ray-irradiated mice, with a significant difference for WB-irradiated group at 2Gy/min, whereas no disparity was noted after proton exposure. Chongsathidkiet *et al*. 2018 revealed that T cells were sequestered in the bone marrow in treatment-naïve GBM patients and GBM mice, thereby inducing lymphopenia (CD4<411 cells/µL vs. 962 cells/µL in controls)^17^. Data likewise demonstrated a splenic contraction in patients (32% mean size reduction), accompanied by a significant decrease in T cell counts. Could brain irradiation induce a secretion of tissue or blood factors that induce a sequestration of T cells in the marrow leading to a peripheral lymphopenia? In a tumoral context, it may be considered as a ‘tumor-imposed mode’^17^, limiting the antitumor lymphocyte capacities and thereby explaining the low efficacy of immunotherapies.

It is known that the spleen can be used as a radiation target in order to evaluate bystander effects^33^. El-Din *et al*. 2017 investigated the abscopal effect after x-ray treatment, demonstrating an increase in p53 and caspase-3^30^, along with DNA damage induction within the spleen in cranially and whole-body irradiated rats. This adds to the evidence that irradiation’s abscopal effect impacts. It also supports the hypothesis of soluble factors transmitted from cerebral irradiated tissues through the blood to peripheral hematopoietic organs. One of the explanations at the origin of a greater lymphopenia to x-rays could be accounted for by direct radio-sensitivity.

*In vitro* data from isolated splenic mice lymphocytes demonstrated that CD3^+^ are biologically more sensitive to protons than X-rays. These observations are consistent with the literature^24,34^. *Ex vivo* irradiation of human peripheral blood lymphocytes with protons resulted in a significantly high number of necrotic cells compared to x-rays’ short-term after irradiation. Protons are likely more efficient in cell-killing, which is most likely due to the production of more irreparable lesions compared with x-rays. This greater radio-sensitivity of lymphocytes to protons cannot be explained our *in vivo* results. Recently, it was postulated that charged particles are likely more immunogenic compared with x-rays, distinctly affecting cell death pathways and leading to increased immunogenicity, thereby sparing more naïve and memory T cells, which is essential to direct and sustain tumor-specific immune responses^35^. This finding may have implications regarding the potential of a given therapeutic modality to produce robust immune modulation via programmed cell death (x-rays) or inflammation (proton therapy). This should stimulate the development of biomarkers pertaining to immune activation, thereby improving proton therapy.

## CONCLUSION

Herein, we demonstrated that proton therapy exerts a conservative effect on leukocytes compared with x-rays. In addition, our work established a link between brain irradiation, peripheral blood compartment immunity, and hematopoietic organs. Taken together, our data likely improve our understanding of hadron therapy effects on inflammation, enabling us to optimize this treatment for patients undergoing brain irradiation therapy, while also suggesting synergistic effects regarding a possible combination with specific immunotherapies.

## Funding & Acknowledgements

This project was co-funded by the Region Normandie, Ligue contre le Cancer and CNRS through the 80ǀPrime program. The authors thank the French National Agency for Research ‘Investissements d’Avenir’ (n°ANR-10-EQPX1401), the “Programme PAUSE–Collège de France and CNRS” and all facilities: *in vivo* irradiation and imaging (Cyceron, CYRCé), cell irradiation (CYCLHAD), animal care (CURB, Oncomodels), histology (VIRTUAL’HIS), and flow cytometry (Normandie Equine Vallee platform, LABEO BIOTARGEN).

## REFERENCES

1. Chou C, Li MO. Tissue-Resident Lymphocytes Across Innate and Adaptive Lineages. Front Immunol. 2018;9:2104.

2. Kansler ER, Li MO. Innate lymphocytes-lineage, localization and timing of differentiation. Cell Mol Immunol. 2019;16(7):627–633.

3. Oble DA, Loewe R, Yu P, Mihm MC. Focus on TILs: prognostic significance of tumor infiltrating lymphocytes in human melanoma. Cancer Immun. 2009;9:3.

4. Fridlender ZG, Sun J, Kim S, et al. Polarization of tumor-associated neutrophil phenotype by TGF-beta: “N1” versus “N2” TAN. Cancer Cell. 2009;16(3):183–194.

5. Geissmann F, Manz MG, Jung S, Sieweke MH, Merad M, Ley K. Development of monocytes, macrophages, and dendritic cells. Science. 2010;327(5966):656-661.

6. Sica A, Larghi P, Mancino A, et al. Macrophage polarization in tumour progression. Seminars in Cancer Biology. 2008;18(5):349–355.

7. Alifieris C, Trafalis DT. Glioblastoma multiforme: Pathogenesis and treatment. Pharmacol Ther. 2015;152:63–82.

8. Han S, Zhang C, Li Q, et al. Tumour-infiltrating CD4(+) and CD8(+) lymphocytes as predictors of clinical outcome in glioma. Br J Cancer. 2014;110(10):2560–2568.

9. Salgado R, Denkert C, Demaria S, et al. The evaluation of tumor-infiltrating lymphocytes (TILs) in breast cancer: recommendations by an International TILs Working Group 2014. Ann Oncol. 2015;26(2):259–271.

10. Klemm F, Maas RR, Bowman RL, et al. Interrogation of the Microenvironmental Landscape in Brain Tumors Reveals Disease-Specific Alterations of Immune Cells. Cell. 2020;181(7):1643–1660.e17.

11. van der Woude LL, Gorris MAJ, Halilovic A, Figdor CG, de Vries IJM. Migrating into the Tumor: a Roadmap for T Cells. Trends Cancer. 2017;3(11):797–808.

12. Tomaszewski W, Sanchez-Perez L, Gajewski TF, Sampson JH. Brain Tumor Microenvironment and Host State: Implications for Immunotherapy. Clin Cancer Res. 2019;25(14):4202–4210.

13. Osipov A, Lim SJ, Popovic A, et al. Tumor Mutational Burden, Toxicity, and Response of Immune Checkpoint Inhibitors Targeting PD(L)1, CTLA-4, and Combination: A Meta-regression Analysis. Clin Cancer Res. 2020;26(18):4842–4851.

14. Yang F, He Z, Duan H, et al. Synergistic immunotherapy of glioblastoma by dual targeting of IL-6 and CD40. Nat Commun. 2021;12:3424.

15. Gabrusiewicz K, Rodriguez B, Wei J, et al. Glioblastoma-infiltrated innate immune cells resemble M0 macrophage phenotype. JCI Insight. 2016;1(2).

16. Prosniak M, Harshyne LA, Andrews DW, et al. Glioma grade is associated with the accumulation and activity of cells bearing M2 monocyte markers. Clin Cancer Res. 2013;19(14):3776–3786.

17. Chongsathidkiet P, Jackson C, Koyama S, et al. Sequestration of T cells in bone marrow in the setting of glioblastoma and other intracranial tumors. Nat Med. 2018;24(9):1459–1468.

18. Nakamura N, Kusunoki Y, Akiyama M. Radiosensitivity of CD4 or CD8 positive human T-lymphocytes by an in vitro colony formation assay. Radiat Res. 1990;123(2):224–227.

19. Grossman SA, Ye X, Lesser G, et al. Immunosuppression in patients with high-grade gliomas treated with radiation and temozolomide. Clin Cancer Res. 2011;17(16):5473–5480.

20. Yovino S, Kleinberg L, Grossman SA, Narayanan M, Ford E. The etiology of treatment-related lymphopenia in patients with malignant gliomas: modeling radiation dose to circulating lymphocytes explains clinical observations and suggests methods of modifying the impact of radiation on immune cells. Cancer Invest. 2013;31(2):140–144.

21. Mendez JS, Govindan A, Leong J, Gao F, Huang J, Campian JL. Association between treatment-related lymphopenia and overall survival in elderly patients with newly diagnosed glioblastoma. J Neurooncol. 2016;127(2):329–335.

22. Hammi A, Paganetti H, Grassberger C. 4D blood flow model for dose calculation to circulating blood and lymphocytes. Phys Med Biol. 2020;65(5):055008.

23. Mohan R, Liu AY, Brown PD, et al. Proton Therapy Reduces the Likelihood of High-Grade Radiation-Induced Lymphopenia in Glioblastoma Patients: Phase II Randomized Study of Protons vs. Photons. Neuro Oncol. Published online August 5, 2020.

24. Miszczyk J, Rawojć K. Effects of culturing technique on human peripheral blood lymphocytes response to proton and X-ray radiation. Int J Radiat Biol. 2020;96(4):424–433.

25. Miszczyk J. Investigation of DNA Damage and Cell-Cycle Distribution in Human Peripheral Blood Lymphocytes under Exposure to High Doses of Proton Radiotherapy. Biology (Basel). 2021;10(2):111.

26. Grosjean C, Quessada J, Nozais M, Loosveld M, Payet-Bornet D, Mionnet C. Isolation and enrichment of mouse splenic T cells for ex vivo and in vivo T cell receptor stimulation assays. STAR Protoc. 2021;2(4):100961.

27. Stone HB, Peters LJ, Milas L. Effect of host immune capability on radiocurability and subsequent transplantability of a murine fibrosarcoma. J Natl Cancer Inst. 1979;63(5):1229–1235.

28. Pham TN, Coupey J, Candeias SM, Ivanova V, Valable S, Thariat J. Beyond lymphopenia, unraveling radiation-induced leucocyte subpopulation kinetics and mechanisms through modeling approaches. J Exp Clin Cancer Res. 2023;42(1):50.

29. Verhaegen F, Butterworth KT, Chalmers AJ, et al. Roadmap for Precision preclinical x-ray radiation studies. Phys Med Biol. Published online December 30, 2022.

30. Mohye El-Din AA, Abdelrazzak AB, Ahmed MT, El-missiry MA. Radiation induced bystander effects in the spleen of cranially-irradiated rats. Br J Radiol. 2017;90(1080):20170278.

31. Suckert T, Beyreuther E, Müller J, et al. Late Side Effects in Normal Mouse Brain Tissue After Proton Irradiation. Front Oncol. 2020;10:598360.

32. Cesaire M, Le Mauff B, Rambeau A, Toutirais O, Thariat J. [Mechanisms of radiation-induced lymphopenia and therapeutic impact]. Bull Cancer. 2020;107(7-8):813–822.

33. Koturbash I, Loree J, Kutanzi K, Koganow C, Pogribny I, Kovalchuk O. In vivo bystander effect: cranial X-irradiation leads to elevated DNA damage, altered cellular proliferation and apoptosis, and increased p53 levels in shielded spleen. Int J Radiat Oncol Biol Phys. 2008;70(2):554–562.

34. Benderitter M, Durand V, Caux C, Voisin P. Clearance of radiation-induced apoptotic lymphocytes: ex vivo studies and an in vitro co-culture model. Radiat Res. 2002;158(4):464–474.

35. Durante M. New challenges in high-energy particle radiobiology. Br J Radiol. 2014;87(1035):20130626.

