## Supplementary material for "Investigating the effects of protons versus x-rays on radiation-induced lymphopenia after brain irradiation": Supplemantary data

| Antibodies | Cellular subtypes | Fluo | Reference | Supplier | Concentration |
|---|---|---|---|---|---|
| **CD16/CD32** | Fcγ receptors | - | 101320 | Biolegend™ | 0.5 mg/ml |
| **CD45.2** | Haematopoietic stem cells | BV650 | 109836 | Biolegend™ | 0.2 mg/ml |
| **CD3** | T lymphocyte | BV605 | 100237 | Biolegend™ | 100 μg/ml |
| **CD4** | Helper T cell | PE | 12004282 | Invitrogen | 0.2 mg/ml |
| **CD8** | Cytotoxic T cell | PE/Cy7 | 100722 | Biolegend™ | 0.2 mg/ml |
| **B220 (CD45.R)** | B lymphocyte | Percp/Cy5.5 | 103236 | Biolegend™ | 0.2 mg/ml |
| **NK1.1** | Natural killer | FITC | 11594182 | Invitrogen | 0.5 mg/ml |
| **RORγt** | Helper T cell 17 (Th17) | PE/TR | 61698182 | Invitrogen | 0.2 mg/ml |
| **FoxP3** | Regulatory T cell | BV421 | 48577382 | Invitrogen | 0.2 mg/ml |
| **CD44** | Effector T cell | FITC | 11044182 | Invitrogen | 0.5 mg/ml |
| **CD62L** | Effector memory T cell | Percp/Cy5.5 | 45062182 | Invitrogen | 0.2 mg/ml |
| **PD-1 (CD279)** | Exhaustion marker | PE | 135206 | Biolegend™ | 0.2 mg/ml |
| **CTLA4 (CD152)** | Exhaustion marker | BV421 | 106312 | Biolegend™ | 0.2 mg/ml |
| **CD11b** | Myeloid | Percp/Cy5.5 | 550993 | BD Biosciences™ | 0.2 mg/ml |
| **SiglecF** | Eosinophils | PE/TR | 130112172 | Miltenyi Biotec™ | 1.5 μg/ml |
| **Ly6C** | Monocyte | BV605 | 563011 | BD Biosciences™ | 0.2 mg/ml |
| **Ly6G** | Neutrophils | BV421 | 127628 | Biolegend™ | 0.2 mg/ml |
| **MHCII** | Monocyte subtype | PE/Cy7 | 107630 | Biolegend™ | 0.2 mg/ml |
| **CX3CR1** | Monocyte subtype | PE | 149006 | Biolegend™ | 0.2 mg/ml |
| **F4/80** | Macrophage | FITC | 11480182 | Invitrogen | 0.5 mg/ml |

**Supplementary table 1. Flow cytometry antibodies**

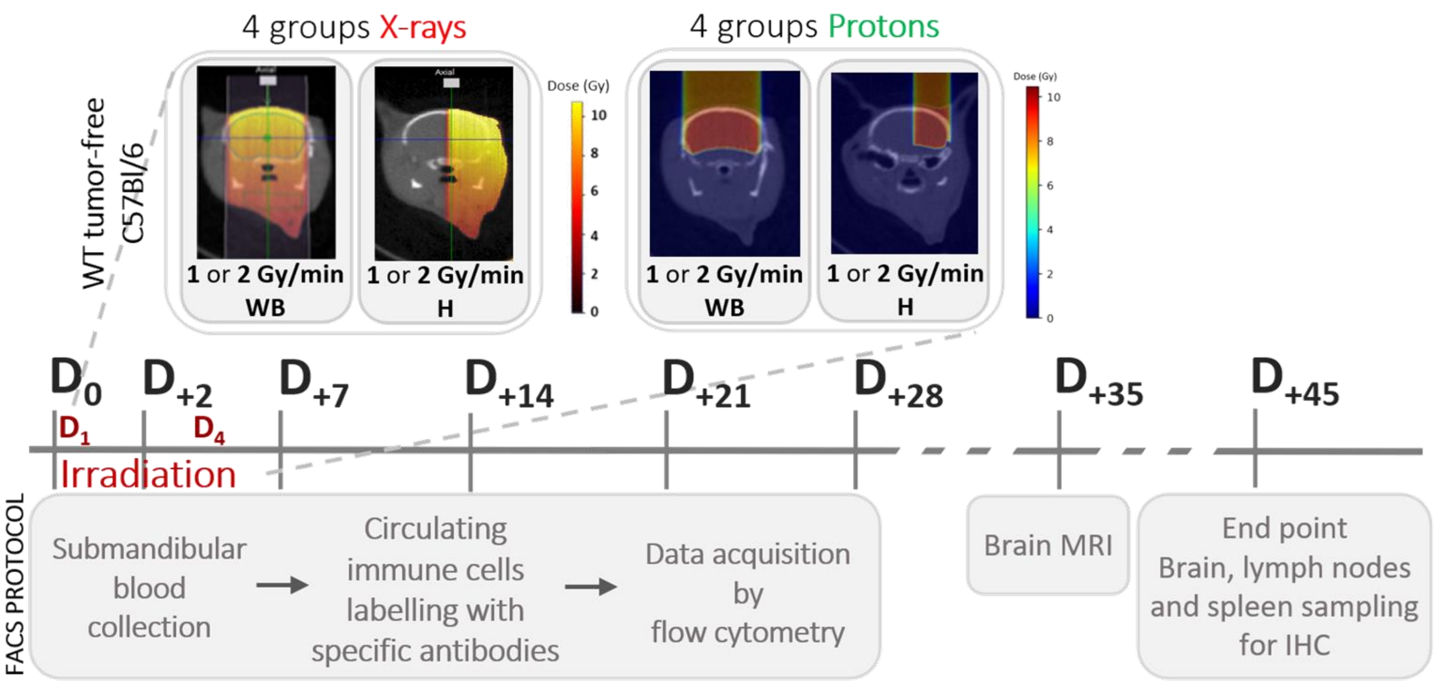


**Supplementary figure 1.** ***In vivo* methods**. ***Experimental protocol of comparative effects of x-ray versys proton by whole peripheral blood collection and circulating immune cells analyses before ($D_0$), during ($D_2$) and after ($D_7$, $D_{14}$, $D_{21}$, $D_{28}$) tumor-free C57BL/6 mice brain irradiation***. Effects on circulating leukocytes were studied using flow cytometry. Whole peripheral blood was collected at six time points ($D_0$, $D_2$, $D_7$, $D_{14}$, $D_{21}$ and $D_{28}$) and labeled with specific antibodies in order to detect and analyze each leukocyte subpopulation. For both x-ray or proton beams, tumor-free C57BL/6 mice were irradiated in 2.5Gy twice-daily sessions for four consecutive days ($D_1$ to $D_4$) according to two variables: irradiation volume (whole-brain (**WB**) or hemisphere (**H**)) and dose rates (1 or 2 Gy/min). At late time, brain MRI was performed in order to check anatomical changes. The protocol end point is marked by the removal of specific organs for subsequent immunohistochemistry (IHC) analyses.

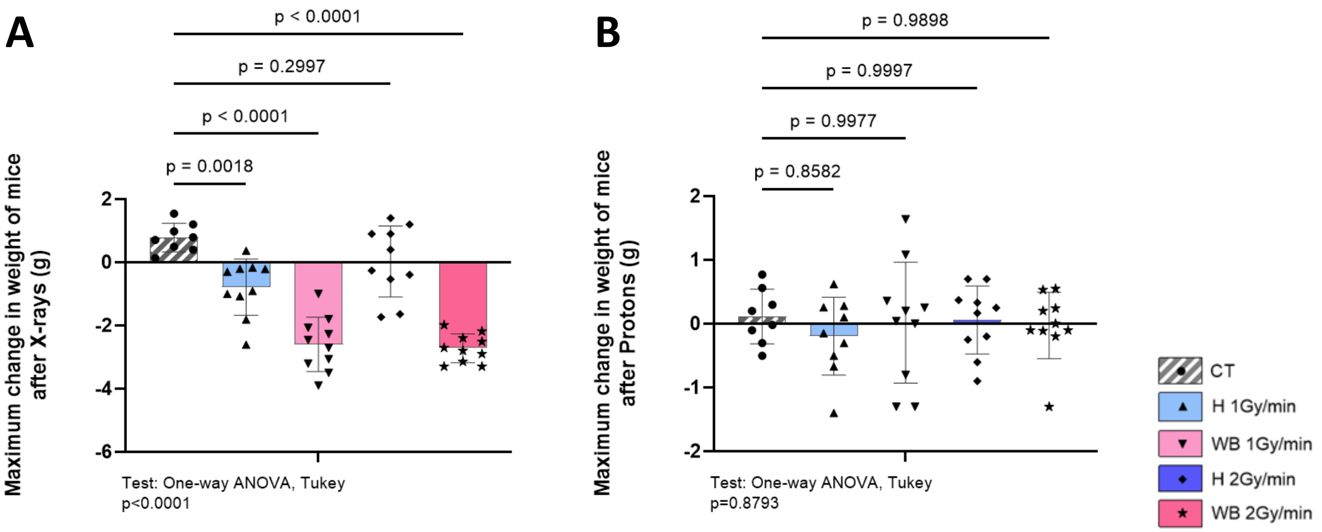


**Supplementary figure 2.** **Maximum change in mice weight occured 10 days after x-ray or proton brain irradiation.** ***Monitoring mice's condition by measuring weight change after x-ray [A] and proton [B] beams (n=10 per groups) compared to controls (CT) (n=8).*** All data were expressed as mean ± standard deviation. P-values were determined using a one-way ANOVA test with a multiple comparison Tukey's test. The most significant negative variation was observed 10 days after X-ray for WB-irradiated groups. The variation after proton brain irradiation followed the controls, being almost zero.

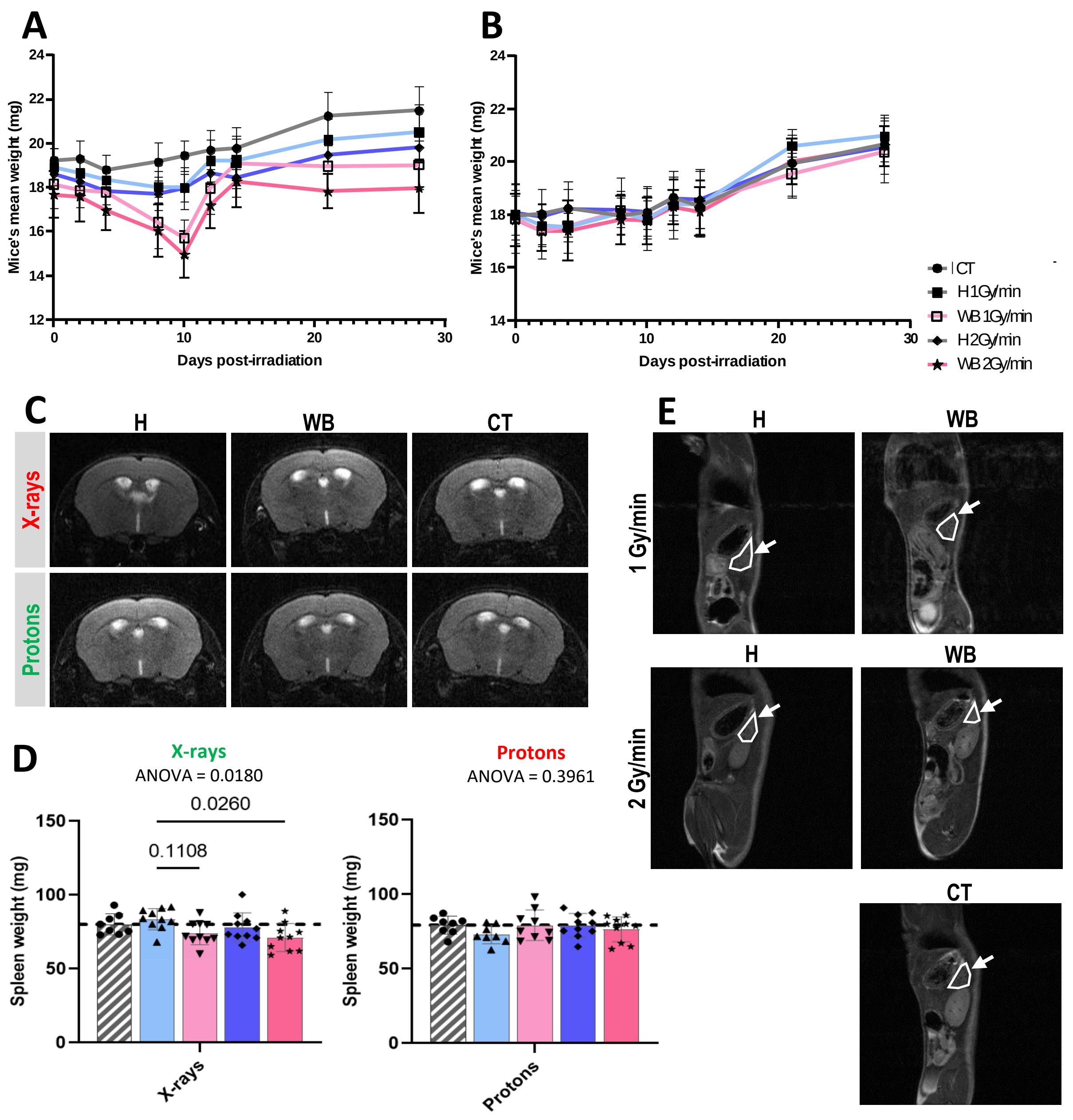


**Supplementary figure 3.** **Crosstalk: brain irradiation, blood compartment and peripheral hematological organs. [A-B]** ***Evolution of mice's mean weight over time after x-rays (A) and protons (B).*** All data were expressed as mean ± standard deviation. **[C]** ***Mice's brain MRI imaging.*** MRI was performed in order to check for macro-anatomical and radio-necrosis changes late ($D_{35}$) after x-rays or protons. For each modality, three animals per group (H or WB at 2Gy/min) and one control (CT) for both x-rays and protons cohort were evaluated. **[D]** ***Spleen weight after x-ray or proton brain irradiation*****.** The histograms represent the spleen weight at the protocol end point ($D_{45}$) for all irradiated mice (n=10 per groups) compared with CT (n=8) after x-rays (*left*) or protons (*right*). All data were expressed as mean ± standard deviation. *P*-values were determined using a one-way ANOVA test, along with a multiple comparison Kruskal-Wallis test. **[E]** ***Mice's whole-body MRI imaging*****.** Whole-body acquisitions were performed on CT, as well as 2Gy/min- and 1Gy/min-irradiated mice late after hemispheric (H) or whole-brain (WB) x-ray irradiation ($D_{35}$). White arrows show the spleen bounded by a white line.

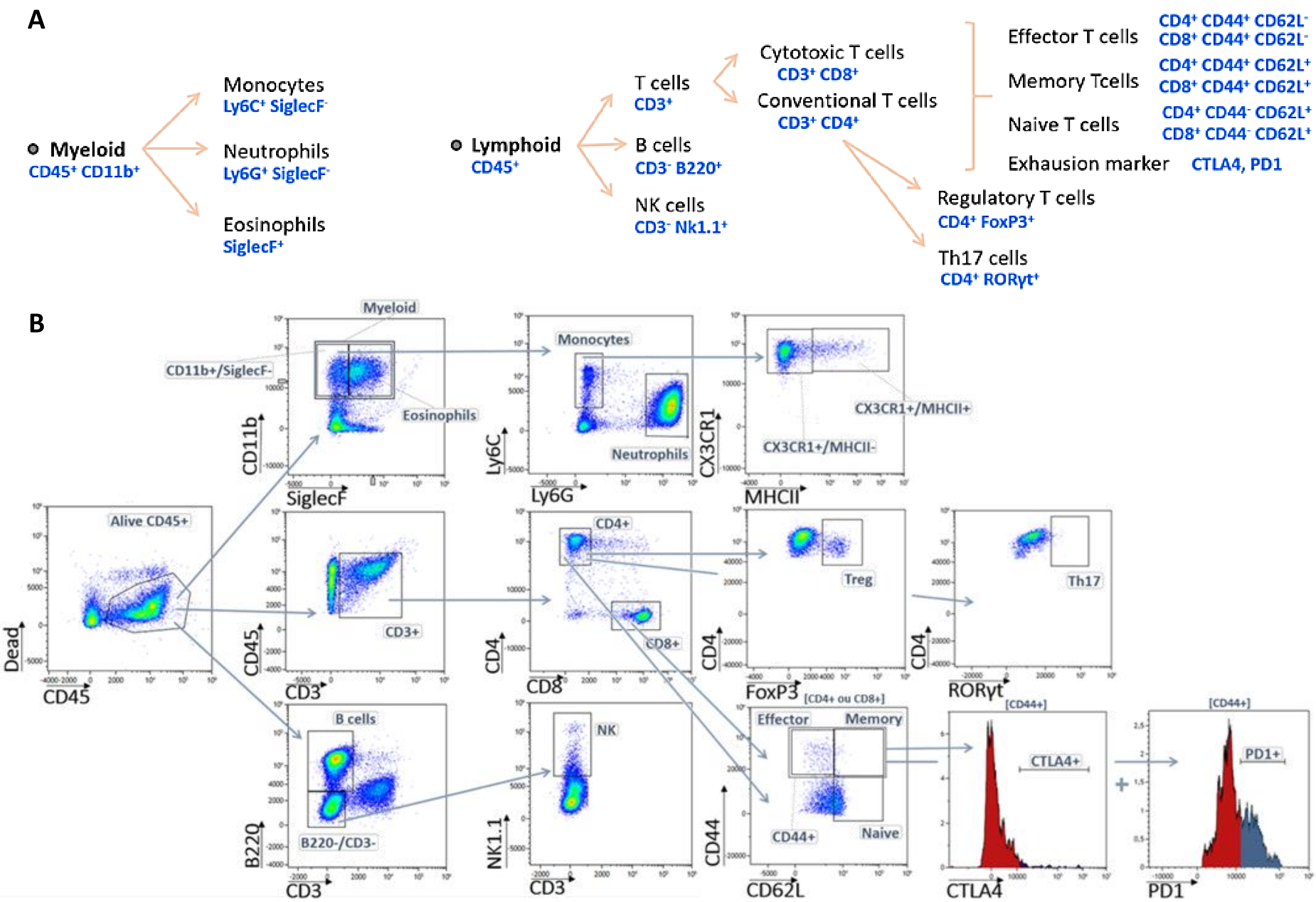


**Supplementary method 1.** **<u>Analyses of preclinical cytometry data</u>. [A]** ***Schematic representation of all circulating leukocyte subtypes of interest***. Specific markers are identified by the CytoFLEX S® allowing for subsequent quantifications. **[B]** ***Gating strategy***. From whole peripheral blood, alive hematopoietic stem cells (dead$^-$CD45$^+$) were selected to gate on leukocyte's lineages: myeloid (CD11b$^+$), T (CD3$^+$), B (CD3$^-$B220$^+$), NK (CD3$^-$NK1.1$^+$) cells allowing for the subpopulation dosage to be refined. From myeloid cells, specific subtypes were identified: monocyte (CD11b$^+$SiglecF$^-$Ly6C$^+$), neutrophil (CD11b$^+$SiglecF$^-$Ly6G$^+$), eosinophil (CD11b$^+$SiglecF$^+$). From T lymphocyte (CD45$^+$CD3$^+$), specific subpopulations were selected: cytotoxic (CD8$^+$), conventional (CD4$^+$), regulatory (CD4$^+$FoxP3$^+$) and helper 17 (CD4$^+$RORγt$^+$) T cells. From CD4$^+$ and CD8$^+$, effector (CD4$^+$CD44$^+$, CD8$^+$CD44$^+$), memory effector (CD4$^+$CD44$^+$CD62L$^+$, CD8$^+$CD44$^+$CD62L$^+$) and naive (CD4$^+$CD44$^-$CD62L$^+$, CD8$^+$CD44$^-$CD62L$^+$) T cells were identified. Subtypes with exhaustion markers (CTLA4, PD1) were detected.
